# Bending and Squeezing: Gradual Potentials Encode Mechanical Stimuli in Poplar

**DOI:** 10.1101/2025.08.25.672115

**Authors:** Erwan Tinturier, Éric Badel, Nathalie Leblanc-Fournier, Geoffroy Guéna, Yoël Forterre, Jean-Louis Julien

**Author notes:** Corresponding authors: *Erwan Tinturier and Jean-Louis Julien Email:.

## Abstract

Mechanical stimuli such as wind elicit rapid electrical signals in plants, yet the mechanisms driving these responses remain poorly understood. Here, we investigated the electrophysiological responses of young poplar trees to controlled stem bending. We identified a gradual potential (GP) distinct from classical action potentials, whose amplitude attenuation and propagation distance were strongly influenced by stimulus speed and intensity. Although maximal GP amplitude near the bending site remained stable across stimulation setups, slower and gentler flexions led to more rapid spatial decay and reduced signal propagation. Similar GP responses were observed following either stem bending or direct root pressurization, suggesting hydraulic-electrical coupling. These results suggest that mechanical stress induces a transient hydraulic pressure wave that triggers the GP. While the GP’s attenuation and propagation dynamics resemble those of a diffusive pressure signal, key features—such as constant peak amplitude regardless of stimulus strength and progressive signal narrowing—point to a complex, non-linear transduction mechanism. Altogether, our findings reveal a finely tuned system where spatial and temporal dynamics of this non-action potential electrical signal encode detailed information about mechanical stimuli, potentially allowing plants to adaptively respond to fluctuating environmental forces like wind.

**Highlight:** Root pressurization generates electrical signals similar to stem bending in poplar, suggesting hydroelectric coupling in plant mechanosensing and wind intensity encoding.

## Introduction

Trees are sessile organisms that grow vertically, gradually developing aerial and underground organs to capture the light, water, and nutrients required for their functioning. These structures can live and grow—sometimes for centuries—in a fluctuating and restrictive environment that exposes them to a range of stimuli. To cope with these challenges, plants have evolved acclimatization mechanisms that allow rapid responses to environmental changes and stresses (Kleine et al., 2021; Zhang et al., 2021).

Environmental stimuli or modulations can be perceived either throughout the entire plant or in specific tissues. When perception is localized, the affected tissue or organ can acclimatize and initiate signaling via the vascular system, triggering responses in distant parts of the plant (Fichman & Mittler, 2021; Li et al., 2021; Suda & Toyota, 2022). This communication between organs involves a variety of mobile signals, including amino acids (Bellandi et al., 2022), peptides (e.g. CLE25; Takahashi et al., 2018, 2019), proteins, miRNAs, hormones, calcium waves, reactive oxygen species (ROS) (Peláez-Vico et al., 2022), hydraulic signals (Malone, 1993; Christmann et al., 2013; Lopez et al., 2014; Louf et al., 2017; Grenzi et al., 2023; Gao et al., 2023; Bacheva et al., 2025), and electrical signals (Pickard, 1973; Wildon et al., 1992; Mousavi et al., 2013; Scherzer et al., 2022; Gao et al., 2023). The integration and coordination of these signaling mechanisms enable plants to adapt to their environment in sophisticated and efficient ways.

Under natural conditions, wind is the primary external force that daily deforms plant organs. The morphological responses to such mechanical stresses are known as thigmomorphogenesis (Jaffe, 1976; Telewski & Jaffe, 1986). Non-wounding mechanical stress, such as organ bending, triggers a local growth response—most notably, an increase in stem diameter in woody species (Leblanc-Fournier et al., 2008). This local thickening response correlates linearly with the cumulative deformation experienced during stress (Coutand et al., 2009). In addition, a distal response affecting elongation growth has been observed (Coutand et al., 2000), which correlates logarithmically with the total applied deformation (Coutand & Moulia, 2000) suggesting the involvement of long-distance signaling mechanisms.

While the morphological responses to non-wounding mechanical stress are well documented, the underlying signaling mechanisms—particularly the interplay between hydraulic and electrical signals—remain much less understood than those involved in wounding responses (Yang et al., 2023). In several woody species, bending of stems and branches generates a hydraulic pressure wave (Lopez et al., 2014; Louf et al., 2017), characterized by a nonlinear poroelastic response in which the pressure pulse varies quadratically with bending curvature and scales with the strained stem volume (Louf et al., 2017). Whether these hydraulic waves are sensed and contribute to long-distance thigmomorphogenetic signaling remains an open question.

Recently, a new type of electrical signal, termed the *Gradual Potential* (GP) has been identified during the bending of poplar and Douglas-fir stems (Tinturier et al., 2021, 2024). This GP propagates rapidly, at a typical velocity of 10–20 cm·s^−1^, and over long distances (up to ∼20 cm). It may represent an autonomous electrical signal, initiated by local cell deformation and propagating electrotonically through local current loops involving excitable cells (Hedrich et al., 2016). Alternatively, the GP may be coupled to the underlying hydraulic pressure wave induced by bending. According to this hydro-electric hypothesis, the GP would arise from deformation of parenchyma cells in contact with xylem vessels as the hydraulic wave propagates, via activation of mechanosensors such as mechanosensitive ion channels (Hamilton et al., 2015; Frachisse et al., 2020).

In this study, we explore the hydro-electric hypothesis for the generation and propagation of the Gradual Potential (GP) induced by stem bending in *Populus tremula × alba*. To this end, we first examine the GP response across a wide range of bending amplitudes and bending rates, as these mechanical parameters are known to strongly influence bending-induced hydraulic waves in stems (Lopez et al., 2014; Louf et al., 2017) as well as electrical potentials triggered by non-damaging touch stimuli (Yang et al., 2023). We then investigate how successive or sustained bending events affect GP generation and identify a refractory period of several hours. Finally, we show that pressurizing the root system and a stem segment to 0.1 MPa—the typical pressure generated during 2% stem bending—elicits a GP with characteristics similar to those observed during bending stimulation. Taken together, these results suggest that mechanically induced hydraulic waves may act as upstream signals for GP generation and for more complex downstream, long-distance signaling responses to non-damaging mechanical stress.

## Materials and methods

### Plant material and culture conditions

Young poplars (*Populus tremula×alba*, clone INRA 717-1B4) were obtained by *in vitro* micropropagation (Leple *et al*., 1992). Once they reached a height of about 4 cm, the acclimation process from *in vitro* to hydroponic solution began (Martin *et al*., 2009) through decreasing relative humidity. Trees were then placed in a growth chamber (16 h/8 h light/dark cycle at 40 μmol m^−2^ s^−1^ and 22°C/18°C with air relative humidity of 60%). Four months after micropropagation, the poplars were used for experiments; at this stage, stems were about 77.8 cm (±1.5 SE) tall with an average diameter of 5.8 mm (±0.3 SE).

Prior to electrophysiology experiments, plants were moved into a Faraday cage in ambient laboratory conditions (16 h/8h light/dark cycle at 20 μmol m^−2^ s^−1^ and 22°C/20°C). Plants were set vertically and fixed in two points of the stem with clamping rings (Fig. 1: Tree support). Foam was rolled around each part of the stem before tightening the clamping rings to avoid stem wounds and to allow possible stem diameter variations. The root system was plunged in a 20 L vat filled with hydroponic solution.

**Figure 1:**
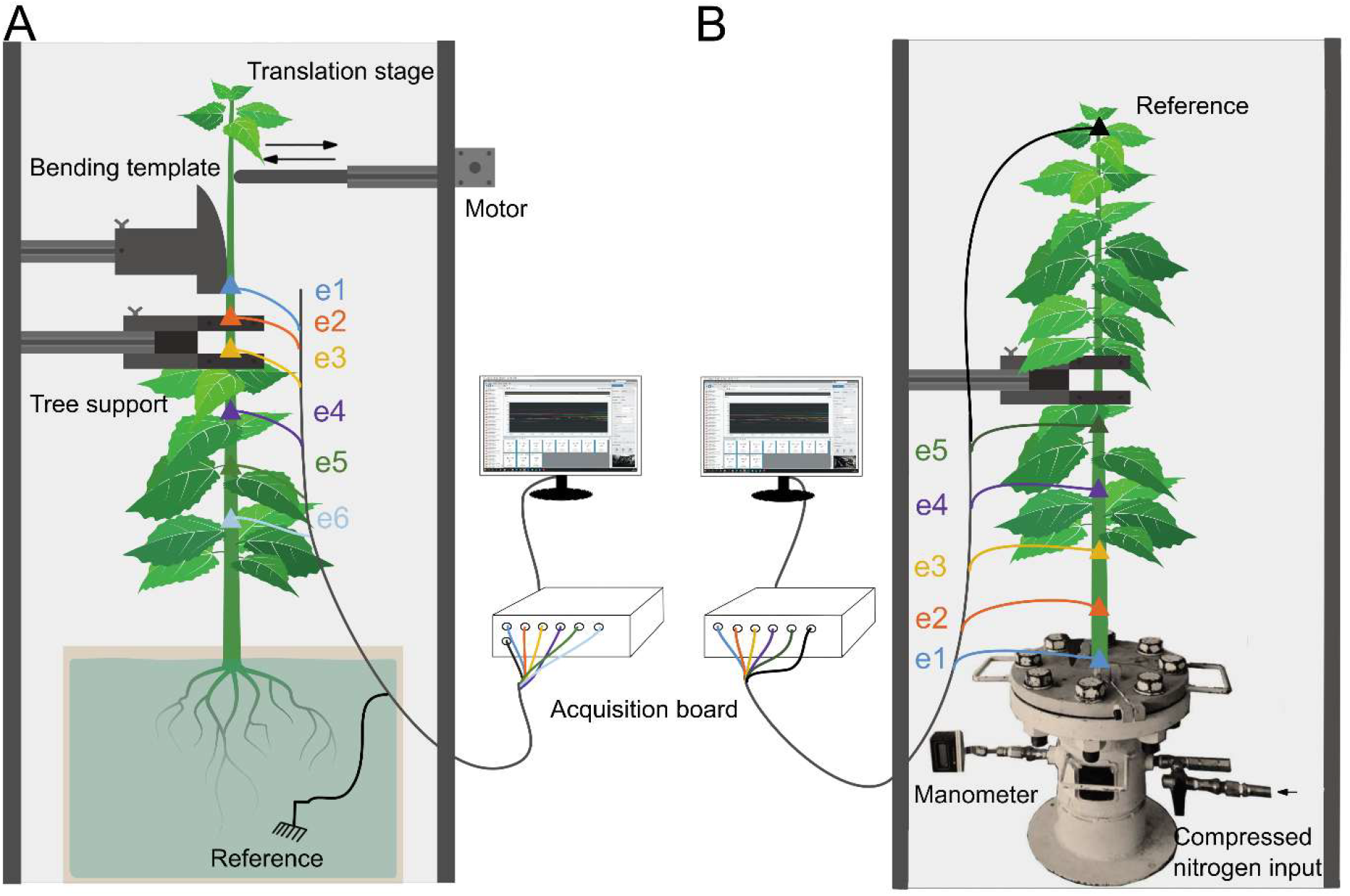
Experimental set-up for bending and pressuring. Poplars were placed in Faraday cages and fixed at two points on the stem using clamping rings (tree support). In the experimental set-up designed for stem bending (A), a plastic circular template (constant radius) was positioned against the stem two centimeters above the tree support. The stem was bent by a motorized arm (motor + translation stage). The electrodes were placed in the immobile portion of the stem. Electrode e1 was inserted 2 cm below the stem-template contact. Subsequently, the distances were as follows: |e1–e2| = 2 cm, |e2–e3| = 3 cm, |e3–e4| = 5 cm, |e4–e5| = 5 cm, and |e5–e6| = 5 cm. The potential difference was measured between each measurement electrode and a reference electrode placed in the bath. In the experimental set-up designed for pressuring (B), the electrodes also were placed in the immobile portion of the stem. Electrode e1 was inserted as close as possible to the outlet of the pressure chamber, at distance 0 cm. Then the other electrodes were inserted relative to e1 as follows: e2 at 5 cm, e3 at 10 cm, e4 at 15 cm, e5 at 22 cm. The potential difference was measured between each measurement electrode and a reference electrode composed of a copper wire inserted at the apex of the plant.

### Bending treatments

The leaves of the bent segment were removed with a razor blade to avoid uncontrolled mechanical stimuli. The speed and the magnitude of the bending stimulation were controlled by a motorized arm (hybrid stepping motors 17PM-H311-P1, Minebea.Co) that pushed the stem against a plastic circular template, which had a constant radius of curvature. The speed “v” of the linear motor was fixed in order to manage three different speed bending: 2.5 cm s^-1^ (fast, 2s forth and back), 0.55 cm s^-1^ (medium) and 0.18 cm s^-1^ (slow) using interface software MegunoLink. The strain magnitude on the stem periphery was controlled by the radius of curvature of the plastic template. This method allowed applying the same strain level along L_0_ = 12 cm of the bent segment. A range of circular plastic templates allowed to adjust the radius of curvature of the template. The radius of curvature was adjusted to apply three different maximal peripheral longitudinal strain of the bark ε_max_ : around 0.2%, 1% and 2 % ; according to the following equation (Coutand *et al*., 2009; Moulia *et al*., 2015).

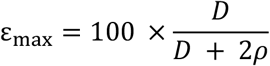

where ρ is the radius of curvature of the plastic template and D is the diameter of the stem in the direction of the bending (Niez *et al*., 2019). This value is high enough to generate significant thigmomorphogenetic responses (Niez *et al*., 2019). We used templates 12 cm long. For each bending experiment, the template was positioned against the stem 35 cm below the apex (Fig. 1). This setup enables to control independently the strain ε_max_ applied on the stem and the strain rate 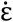; which can be evaluated as 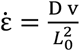. Thanks to the control of the motor speed, we applied three levels of strain rate; 0.07 % s^-1^, 0.2 % s^-1^ and 1 % s^-1^.

### Monitoring of extracellular electrical signals

One single poplar at a time was placed in a Faraday cage. All electronic materials were located outside the cage. Measuring electrodes (tinned copper wire (359-835), 0.25 mm diameter, RS Components) were inserted into the stem, passing through all the tissue to the pith, and measured electrical potential simultaneously near the bending area and up to 20 cm from it. After the installation was completed, the plant was left undisturbed for stabilization for 24 to 48 hours.

The reference electrode (RC3 model, World Precision Instruments) was made of an Ag/AgCl wire immersed in the nutrient solution of the plants. The electrical potential recorded is the difference between a measuring electrode and the reference electrode with respect to the ground. In contrast to intracellular measurements, extracellular measurements show potential changes downward for depolarization and upward for hyperpolarization. For bending experiment, the first measuring electrode e1 was inserted 2 cm below the stem-to-template contact (Fig. 1A). The respective distances of each electrode from e1 were 2 cm (e2), 5 cm (e3), 10 cm (e4), 15 cm (e5), and 20 cm (e6). The measuring electrodes and the reference electrode were connected to a data acquisition card (cDAQ-9171, National Instrument). The card was used as an impedance amplifier (10 GΩ) and A/D converter. DAQExpress 1.0.1 software (National Instrument) monitored the potential difference with a sampling rate of 10 Hz. The graphs and analyzes were processed using MATLAB® software.

The analysis of the recordings provided several parameters of the signal. The amplitude was defined as the maximum difference value compared to the baseline before bending or pressuring. The average propagation speed of the signal was calculated between each electrode as the ratio of the distance between two consecutive electrodes to the delay between the detection of a change in potential difference.

### Pressure treatments and monitoring of extracellular electrical signals

The poplar roots and 10 to 15 cm of stem were enclosed in a 20-cm diameter pressure chamber hermetically separated from the tree aerial part (Fig. 1B). The root system was partially plunged in hydroponic solution (1L) inside the pressure chamber. The pressure chamber is connected to a nitrogen cylinder. A valve allowed the pressure in the chamber to increase around 0.1MPa in 8 seconds, and another valve is used to purge the system. To monitoring electrophysiology measurements, the pressure chamber was installed in the faraday cage.

Similar to the stem bending experiment, the measurement electrodes were inserted into the stem. The first electrode (e1) was inserted as close as possible to the outlet of the pressure chamber, at distance 0 cm. Then the other electrodes were inserted relative to e1 as follows: e2 at 5 cm, e3 at 10 cm, e4 at 15 cm, e5 at 22 cm. The reference electrode was changed to a copper wire to be inserted at the apex of the plant.

### Determination of the maximum propagation distance of the GP

To compare the GP propagation parameters between two experimental conditions (cond.1 vs. cond.2), we used a piecewise linear model with three parameters. The amplitude of the signal as a function of distance *x* from the source was modeled as follows:

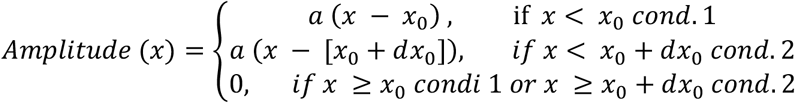

Where:

*a*: slope quantifying the decrease in amplitude with distance, assumed to be identical for both conditions.

*x*_0_: maximum propagation distance in cond.1.

*dx*_0_: difference in maximum propagation distance between cond.1 and cond.2.

The parameter *dx*_0_ was included to account for potential differences in the maximum distance reached by the signal between the two conditions. The slope *a* was constrained to be the same for all conditions. The parameters *a, x*_0_, and *dx*_*0*_ were estimated using non-linear regression via the `nls` function in R software (Team, 2023). Confidence intervals (95%) for each parameter were calculated using the *`confint`* function, allowing for statistical testing of the null hypothesis that *dx*_0_= 0. A significant deviation of *dx*_0_ from zero was interpreted as evidence of a difference in maximum propagation distance between the two conditions.

### Statistical analysis

All measured and computed data were statistically analyzed using R software. Kruskal and Wallis’s tests and Dunn’s tests were performed to compare results in terms of amplitude, duration, and speed of the signal (p < 0.05).

## Results

### Bending amplitude affect GP propagation properties but not initial amplitude

Both electrical and hydraulic signals may depend on the intensity of the mechanical stimulus (bending strain). In plants, the growth response to mechanical stimulus has also been shown to depend on strain, as described by the S3M model (Moulia et al., 2015). We first tested the impact of bending intensity on the characteristics of the Gradual Potential (GP), a key electrical signal observed in response to mechanical deformation. This setup consisted of performing a complete bending cycle (forth and back) of the stem against the template, measured from 35 cm below the apex.

For these experiments, a fixed fast bending speed was applied. This setup corresponds to a high strain rate value 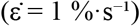. To investigate the effect of bending intensity (ε_max_) on GP characteristics, three bending modalities (ε_max_ = 0.2%, ε_max_ = 1%, and ε_max_ = 2%) were tested.

A GP was induced in all tested modalities (Fig. 2A). The average GP amplitudes measured close to the bending site (0 cm, electrode e1) showed some variability but no clear trend related to strain intensity. Specifically, the mean amplitude at 0 cm was 36.0 ± 1.0 mV for ε_max_ = 2%, 29.7 ± 5.3 mV for ε_max_ = 1%, and 40.6 ± 2.7 mV for ε_max_ = 0.2% (Fig. 2B). As distance from the bending site increased, GP amplitudes generally decreased, with this attenuation being more pronounced at lower bending intensities (Fig. 2B).

**Figure 2:**
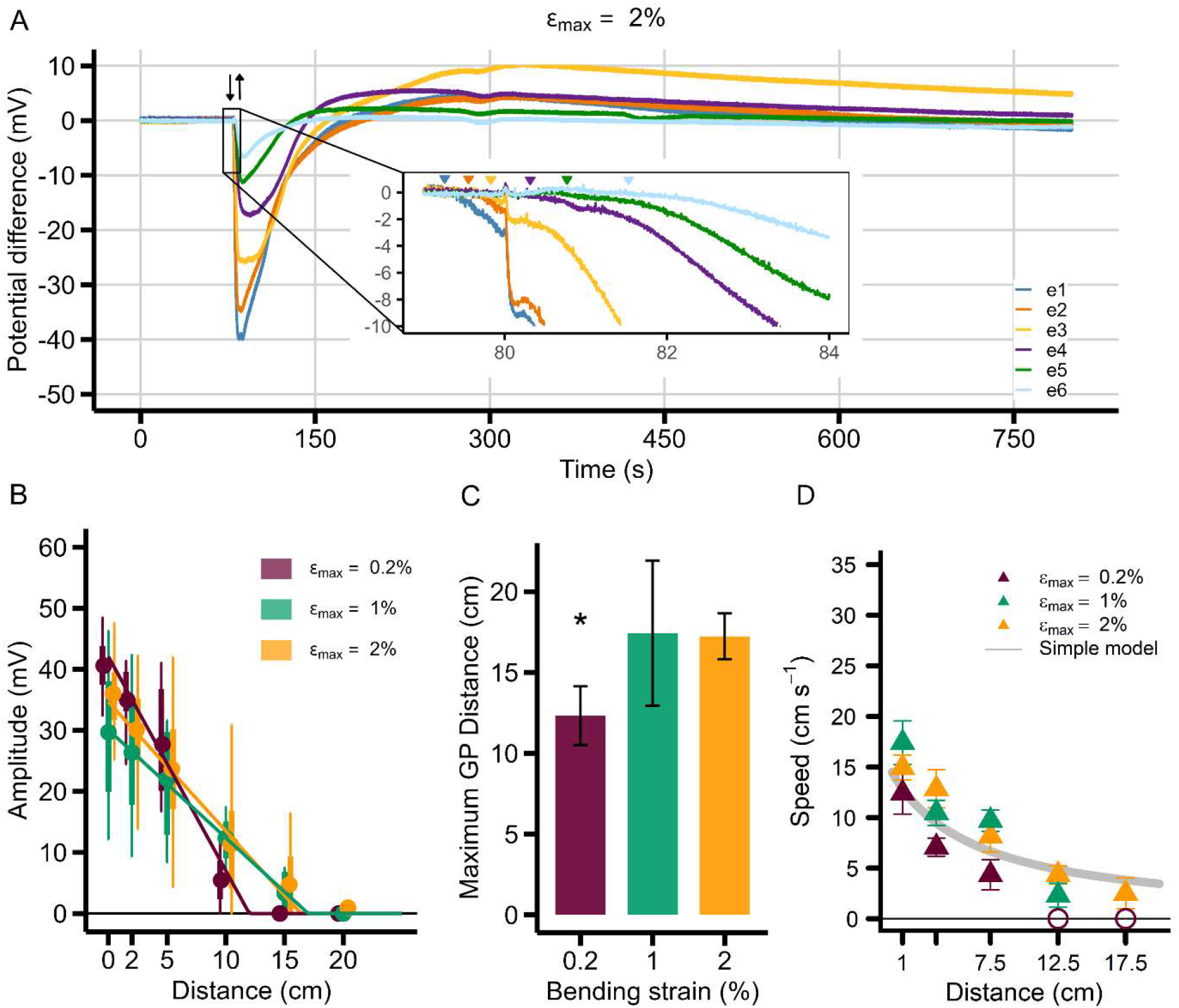
Influence of poplar stem bending intensity (ε) on GP characteristics at a constant bending rate. (A) Representative six-channel potential difference recordings illustrating the GP response to an apical bending with a maximal strain (ε_max_) of 2%. The black arrows indicate the onset (forth) and offset (back) of the bending movement (a complete cycle bending). Electrodes e1 to e6 represent measurements at different distances from the bending site, as described in Figure 1. (B) Boxplot summarizing GP amplitudes as a function of distance from the bending site for three maximal bending intensities: ε_max_ = 0.2% (n=5), ε_max_ = 1% (n=6), and ε_max_ = 2% (n=40). Circles indicate the mean. The solid lines represent the piecewise linear regression fitted to the data for each bending intensity. (C) Estimated maximum propagation distance of the induced GP for the three bending intensities, calculated from piecewise linear regression. The asterisk (*) denotes a significant difference compared to the 2% bending intensity (95% CI). (D) Mean GP propagation velocity as a function of distance (triangles indicate the mean, error bars represent standard error) for the three bending intensities, overlaid with prediction of simple hydraulic diffusive model 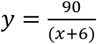 (Simple model, grey line, see Discussion and Supplementary Text S1).

The maximum propagation distance (D_max_), determined from the piecewise linear regressions of amplitude attenuation (Fig. 2B), varied significantly with bending intensity (Fig. 2C). For the 0.2% bending intensity, D_max_ was 12.3 ± 1.8 cm. This value was significantly lower than those observed for the higher bending intensities; 17.4 ± 4.5 cm and 17.2 ± 1.4 cm for 1% and 2% bending intensities respectively, as indicated by the non-overlapping 95% confidence intervals (Fig. 2C). No significant difference in D_max_ was observed between the 1% and 2% bending intensities.

The mean propagation velocity of the GP as a function of distance for the three bending intensities is presented in Fig. 2D. For all intensities, GP propagation velocity consistently decreased hyperbolically with increasing distance from the bending site. Visually, higher bending intensities (1% and 2%) generally resulted in greater propagation velocities, especially at shorter distances, compared to the 0.2% bending intensity. For instance, at 1 cm from the bending site, the mean velocity was 14.9 ± 1.2 cm·s^−1^ for ε_max_ = 2%, 17.4 ± 2.1 cm·s^−1^ for ε_max_ = 1%, and 12.4 ± 2.1 cm·s^−1^ for ε_max_ = 0.2%. At 7.5 cm, these values were 8.2 ± 1.6 cm·s^−1^ (ε_max_ = 2%), 9.7 ± 1.1 cm·s^−1^ (ε_max_ = 1%), and 4.4 ± 1.5 cm·s^−1^ (ε_max_ = 0.2%).

These results indicate that the GP’s capacity for long-distance propagation (reflected in D_max_ and overall propagation velocity profiles) is significantly influenced by the bending strain.

### Bending rate affect GP propagation properties but not initial amplitude

Previous experiments indicated that the signal amplitude does not depend on the bending strain or underlines a saturation effect. Biological sensors sometimes detect the time scale over which a stimulus is applied rather than its amplitude (Maksaev and Haswell 2012). Thus, the time scale of the applied stress is sometimes a relevant parameter. In the next experiments, we varied the bending strain rate for a fixed maximal strain of 2% and a bent stem length of 12 cm.

We examined the effect of bending strain rate 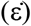 on the GP by performing apical flexions at 2% maximal strain according to three modalities: a low rate 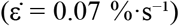, a medium rate 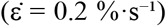, and a high rate 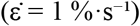. A GP was induced in all tested modalities (Fig. 3A).

**Figure 3:**
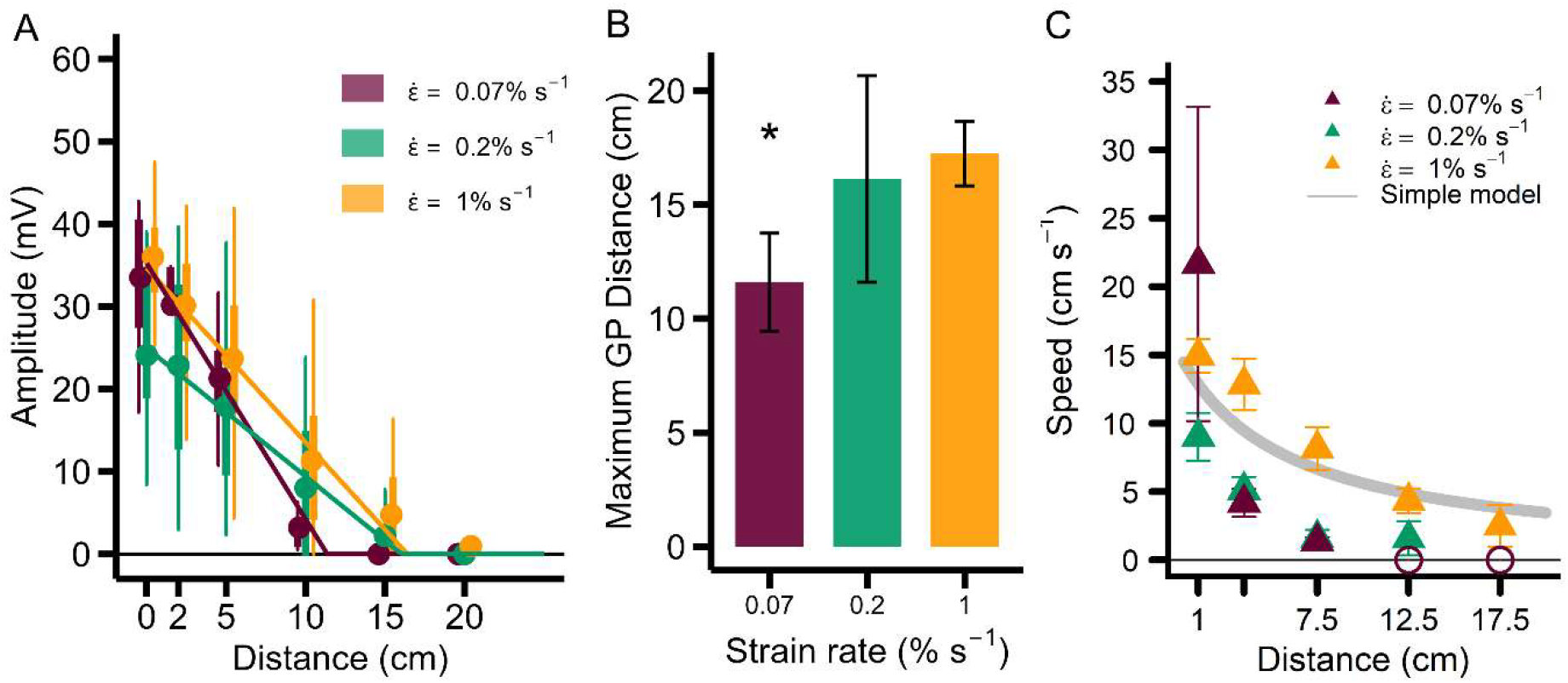
Influence of Poplar Stem Bending Strain Rate 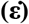 on Gradual Potential (GP) Characteristics. (A) Boxplot summarizing GP amplitudes as a function of distance from the bending site for three bending strain rates: 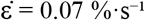 (n=5 plants), 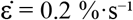 (n=9 plants), and 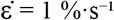 (n=40 plants). Circles indicate the mean. (B) Estimated maximum propagation distance (D_max_) of the induced GP for the three bending strain rates, calculated from piecewise linear regression. The asterisk (*) denotes a significant difference compared to the 1 %·s^−1^ strain rate (95% CI), specifically for 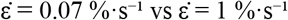. (C) Mean GP propagation velocity as a function of distance (triangles indicate the mean, error bars represent standard error) for the three bending strain rates, overlaid with prediction of simple hydraulic diffusive model 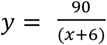 (Simple model, grey line, see Discussion and Supplementary Text S1).

Average GP amplitudes measured at the bending site (0 cm, electrode e1) exhibited variability but no clear monotonic trend with strain rate (Fig. 3A). Specifically, the mean amplitude at 0 cm was 30.1 ± 1.0 mV for 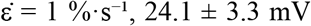 for 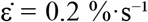, and 33.5 ± 4.9 mV for 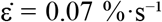. As distance from the bending site increased, GP amplitudes generally decreased, with attenuation appearing more pronounced at lower strain rates. For instance, at 10 cm, mean amplitudes were 11.3 ± 1.2 mV for fast bending, 8.0 ± 3.4 mV for medium bending, and 3.2 ± 1.1 mV for slow bending (Fig. 3A).

The maximum propagation distance (D_max_), estimated using a piecewise linear function fitted to the amplitude data (Fig. 3A), varied significantly with the bending strain rate (Fig. 3B). D_max_ for the slow strain rate 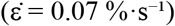 was 11.6 ± 2.2 cm, significantly lower than that for the fast strain rate 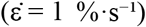, which was 17.2 ± 1.4 cm, as indicated by non-overlapping 95% confidence intervals (Fig. 3B). The medium strain rate 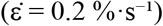 showed an intermediate D_max_ of 16.1 ± 4.5 cm, not significantly different from the fast strain rate.

Mean GP propagation velocity as a function of distance for the three bending strain rates is presented in Figure 3C. For all strain rates, GP propagation velocity consistently decreased hyperbolically with increasing distance (Fig. 3C). Velocity estimates at the shortest distances (e.g., at 1 cm) exhibited larger standard errors (SE) for the slow rate compared to fast and medium rates (Fig. 3C). Mean velocities observed at 1 cm were 14.9 ± 1.2 cm·s^−1^ for 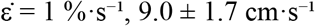 for 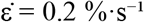, and 21.7 ± 11.5 cm·s^−1^ for 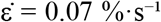. At 7.5 cm, these values were 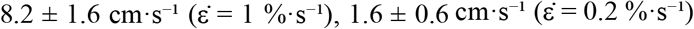, and 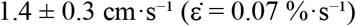.

Once again, the maximum depolarization did not strongly depend on the bending strain rate. However, the propagation length and velocity profiles were significantly modulated by the bending strain rate.

### Role of the bending history

In our reference case experiment, the system is brought back to its initial rest position before the signal is emitted at some distance from the stimulated area. This forthward and backward motion might somewhat impact the overall signal. We tested whether the signal generated by unbending is symetrical to that resulting from bending, or whether it depends on the duration for which the bending strain is maintained. In this attempt, the signal that should be generated during the backward motion may be hindered by the refractory period induced by the preceding forward motion. In the next experiments, we aimed to verify that signal during unbending can occur once the refractory period has elapsed.

### Evidence of a refractory period

We investigated the temporal dynamics of GP refractoriness by applying a second complete bending cycle at various time intervals after an initial bending (Fig. 4A). For every bending, a maximal strain (ε_max_) of 2% and a bending strain rate 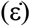 of 1%·s^−1^ were used. The interval between two bendings was 2.5 minutes (2’30), 4 hours (4h), or 7 hours (7h).

**Figure 4:**
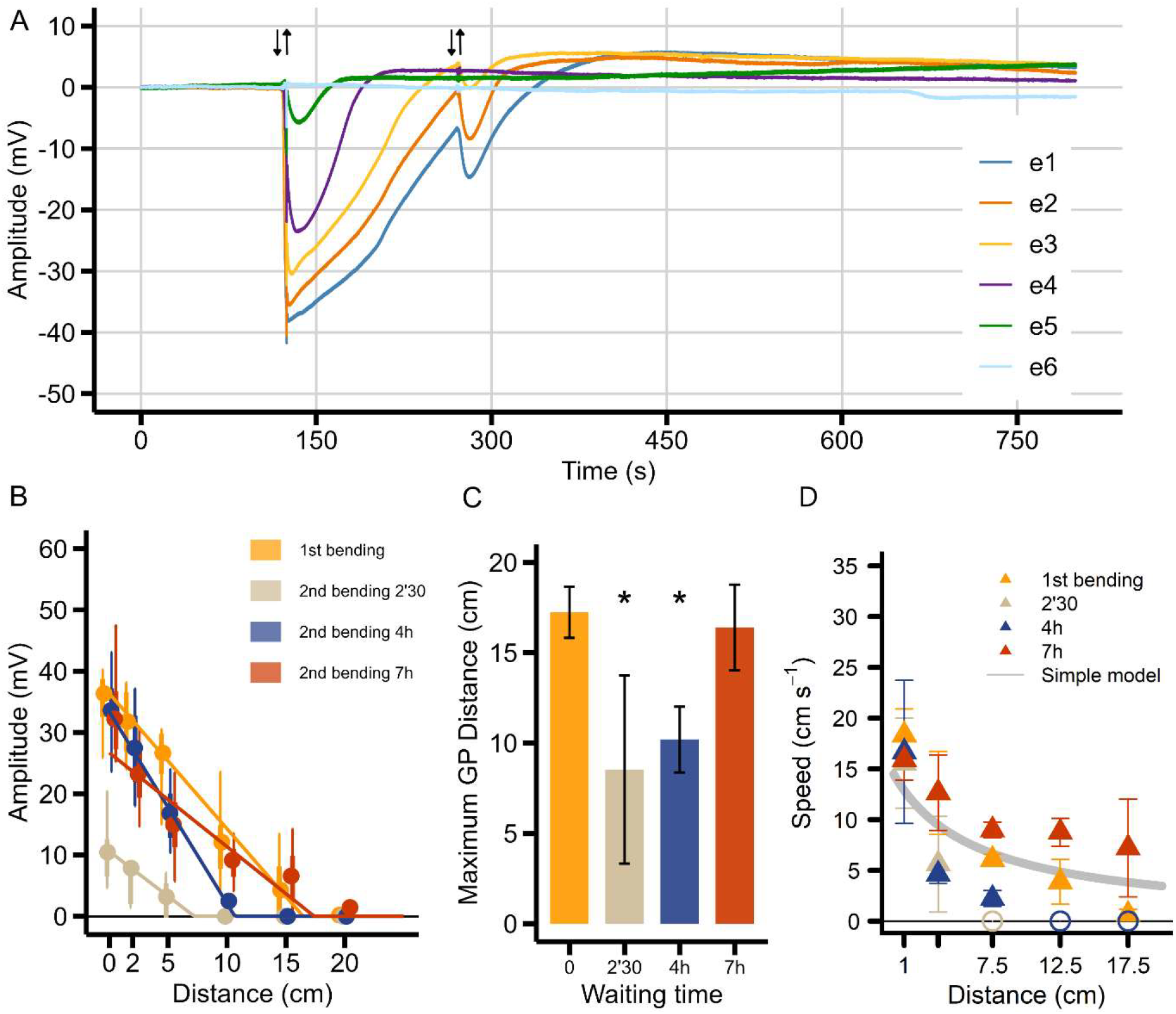
Temporal dynamics of GP refractoriness following a second transient apical bending. (A) Representative recording showing electrical responses at six electrodes following two consecutive bending events separated by 2.5 minutes (the black arrows indicate the onset (forth) and offset (back) of the bending movement (two complete cycles bending). (B) Boxplot summarizing GP amplitudes as a function of distance from the bending site for the initial bending (first Bending) and subsequent bendings applied after 2.5 minutes (2’30), 4 hours (4h), and 7 hours (7h) after the first stimulation (n=5 plants; circles represent the mean). (C) Estimated maximum propagation distance of the GP induced by the first bending and the second bending applied after the three different delay. Maximum distance was estimated using a piecewise linear function; asterisks (*) indicate a significant difference compared to the 1st bending (95% CI, non-overlapping intervals). (D) Mean GP propagation velocity as a function of distance (triangles indicate the mean, error bars represent standard error) for the first bending and the second bendings after 2’30, 4h, and 7h, overlaid with prediction of simple hydraulic diffusive model 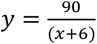 (Simple model, grey line, see Discussion and Supplementary Text S1).

Average GP amplitudes at the bending site (0 cm, electrode e1) showed a significant reduction immediately following the first bending (Fig. 4B). The mean amplitude at 0 cm was 36.4 ± 2.3 mV for the first bending. This amplitude was sharply reduced to 10.5 ± 2.8 mV for the second bending at 2’30, demonstrating a clear refractory period. The amplitude recovered to 33.6 ± 2.7 mV after 4h and 32.2 ± 2.1 mV after 7h, approaching the initial bending amplitude (Fig. 4B). Similarly, GP signal attenuation with distance was more pronounced after 2’30, with amplitudes dropping to baseline much faster compared to the first bending and showing a progressive recovery at 4h and 7h (e.g., at 10 cm, 12.1 ± 2.1 mV for 1st bending, 0 mV after 2’30, 2.5 ± 1.2 mV after 4h, and 9.2 ± 0.8 mV for 7h).

The maximum propagation distance (D_max_), estimated from piecewise linear regressions (Fig. 4B), confirmed these observations (Fig. 4C). The D_max_ for the first bending was 17.2 ± 1.4 cm. It was significantly reduced for the second bending after 2’30 (8.5 ± 5.2 cm) and after 4h (10.2 ± 1.8 cm). However, after 7h, the D_max_ of 16.4 ± 2.4 cm was not significantly different from the first bending, suggesting a full recovery of the propagation capacity.

Mean GP propagation velocity (Fig. 4D) consistently decreased hyperbolically with increasing distance. The immediate refractory period after 2’30 significantly impaired propagation velocity, particularly at longer distances, with velocities dropping quickly to zero. The fastest propagation at 1 cm was observed for the first bending (18.3 ± 2.6 cm·s^−1^), followed by 7h (15.9 ± 2.0 cm·s^−1^), 4h (16.7 ± 7.0 cm·s^−1^), and 2’30 (15.6 ± 4.4 cm·s^−1^). At 7.5 cm, these values were 6.1 ± 0.5 cm·s^−1^ (first bending), 9.0 ± 0.8 cm·s^−1^ (after 7h), 2.2 ± 0.8 cm·s^−1^ (after 4h), and 0 cm·s^−1^ (after 2’30).

These results demonstrate a clear GP refractory period affecting both amplitude and propagation distance/velocity, with a recovery time of approximately 7 hours for full signal propagation capacity.

### Both bending and unbending induce GP

We investigated the electrical response of the stem by separating the bending process into two distinct phases: an initial sustained bending and the subsequent unbending. For the initial bending, a maximal strain (εmax) of 2% and a bending strain rate 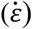 of 1%·s^−1^ were applied. The stem was bent and held for 4 hours (sustained bending) before being returned to its original position (unbending) (Fig. 5A). Both the sustained bending and unbending induced a GP. The “reference bending” (complete cycle from Figures 2-4) is included for comparison in panels B-D.

**Figure 5:**
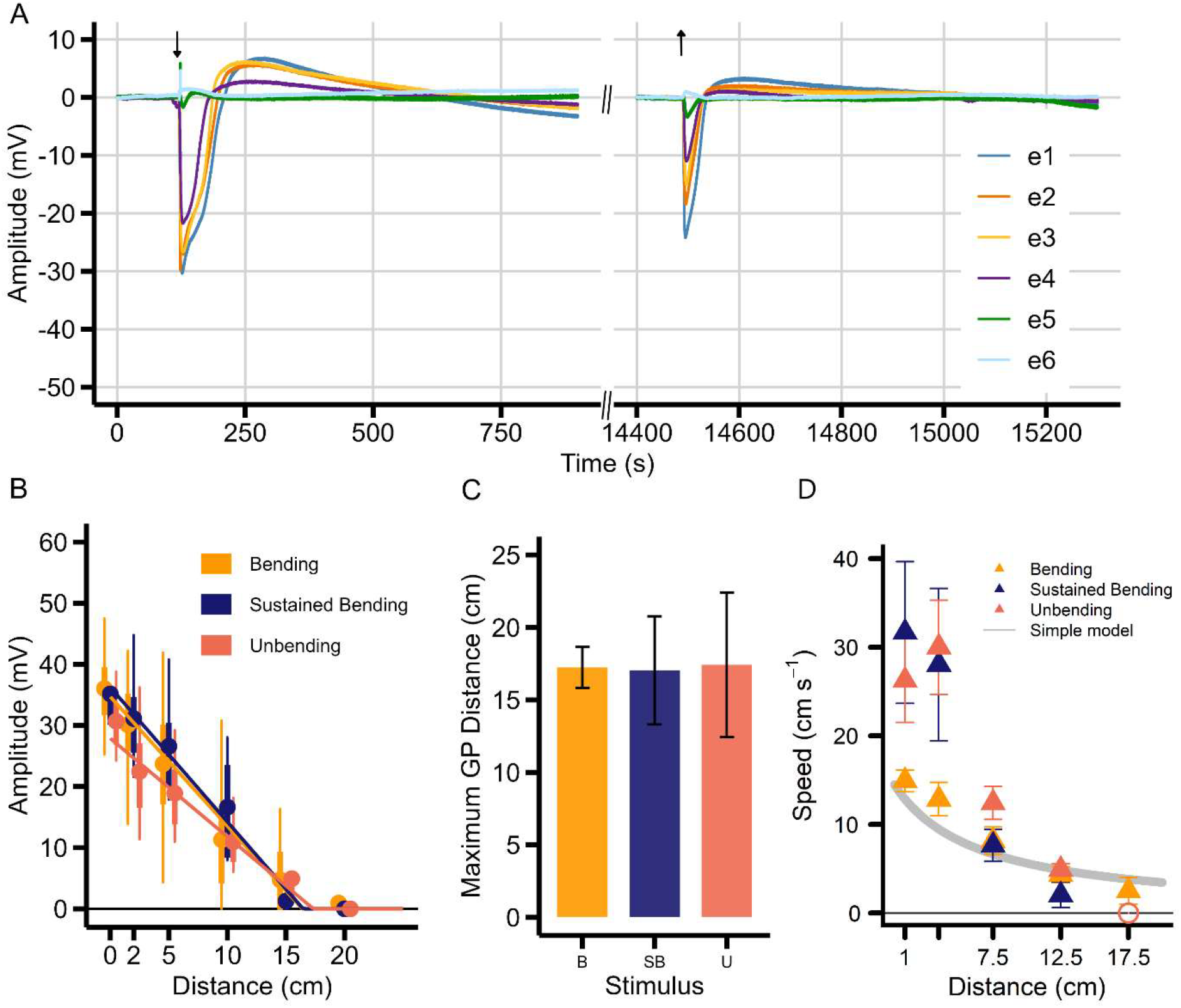
Electrical responses of poplar stem to bending and unbending motions. (A) Representative 6-channel recording illustrating the electrical responses during a sustained bending experiment. The first black arrow indicates the initial bending (start of Sustained Bending) where the stem is held against the template. The second black arrow, 4 hours later, indicates the unbending (return to the initial straight position). (B) Boxplot of GP amplitudes as a function of distance from the bending site for three conditions: “Reference Bending” (complete bending cycle, n=40 plants), “Sustained Bending” (n=4 plants), and “Unbending” (release after 4h sustained bending, n=4 plants). Circles indicate the mean. (C) Estimated maximum propagation distance (D_max_) of the GP induced by “Reference Bending” (B), “Sustained Bending” (SB), and “Unbending” (U), calculated from piecewise linear regression (95% CI). No significant differences were observed between these conditions. (D) Mean GP propagation velocity as a function of distance (triangles indicate the mean, error bars represent standard error) for “Reference Bending”, “Sustained Bending”, and “Unbending”, overlaid with prediction of simple hydraulic diffusive model 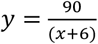 (Simple model, grey line, see Discussion and Supplementary Text S1).

Average GP amplitudes at the bending site (0 cm, electrode e1) showed similar initial depolarizations across all conditions (Fig. 5B). Mean amplitudes were 36.0 ± 1.0 mV for “reference bending”, 35.2 ± 4.4 mV for “sustained bending”, and 30.7 ± 3.1 mV for “unbending”. All conditions exhibited similar GP amplitude attenuation with distance (Fig. 5B). For example, at 10 cm, mean amplitudes were 11.3 ± 1.2 mV for “reference bending”, 16.6 ± 5.0 mV for “sustained bending”, and 10.9 ± 2.6 mV for “unbending”.

Maximum propagation distance (D_max_) showed no significant differences between conditions (Fig. 5C). D_max_ values were 17.2 ± 1.4 cm for “reference bending”, 17.0 ± 3.7 cm for “sustained bending”, and 17.4 ± 5.0 cm for “unbending”, indicating consistent GP propagation length regardless of the bending direction or duration.

Mean GP propagation velocity (Fig. 5D) consistently decreased hyperbolically with distance. Despite visual suggestions of differences, Pairwise Wilcoxon rank-sum tests (Holm correction) revealed no significant differences in GP propagation velocities between bending conditions at any distance interval (all p_adj_ > 0.05). At 1 cm, velocities were 14.9 ± 1.2 cm·s^−1^ (“reference bending”), 31.7 ± 8.0 cm·s^−1^ (“sustained bending”), and 26.3 ± 4.7 cm·s^−1^ (“unbending”). At 7.5 cm, these values were 8.2 ± 1.6 cm·s^−1^ (“reference bending”), 7.6 ± 1.8 cm·s^−1^ (“sustained bending”), and 12.4 ± 1.9 cm·s^−1^ (“unbending”).

These results indicate that sustained bending induces an initial GP comparable to a complete bending cycle, rather than a steady signal. Furthermore, unbending, after a sustained bending, also generates a GP with similar amplitude, propagation distance, and velocity characteristics as the initial bending, provided the refractory period has passed.

### Root pressurization induces a GP with similar characteristics as those of bending

Assuming the hypothesis that bending strain is perceived but may not be the transported signal, and considering that bending generates hydraulic pressure waves, it is important to distinguish between hydraulic pressure and the mechanical stress caused by bending.

To test the hydroelectric coupling hypothesis, we induced a pressure wave in the stem’s vascular system using a hermetically sealed pressure chamber. The root system and a 15 cm stem section were placed in the chamber, with five measuring electrodes positioned at 5 cm intervals along the stem (Figure 1B). A representative recording (Fig. 6A) illustrates this response, which instantaneously affected the first electrode (e1) with a mean amplitude of 28.4 ± 2.9 mV. This response consisted of a rapid depolarization followed by an equally rapid repolarization, lasting about 40 seconds. Notably, no exudation was observed at the leaf level under these conditions.

**Figure 6:**
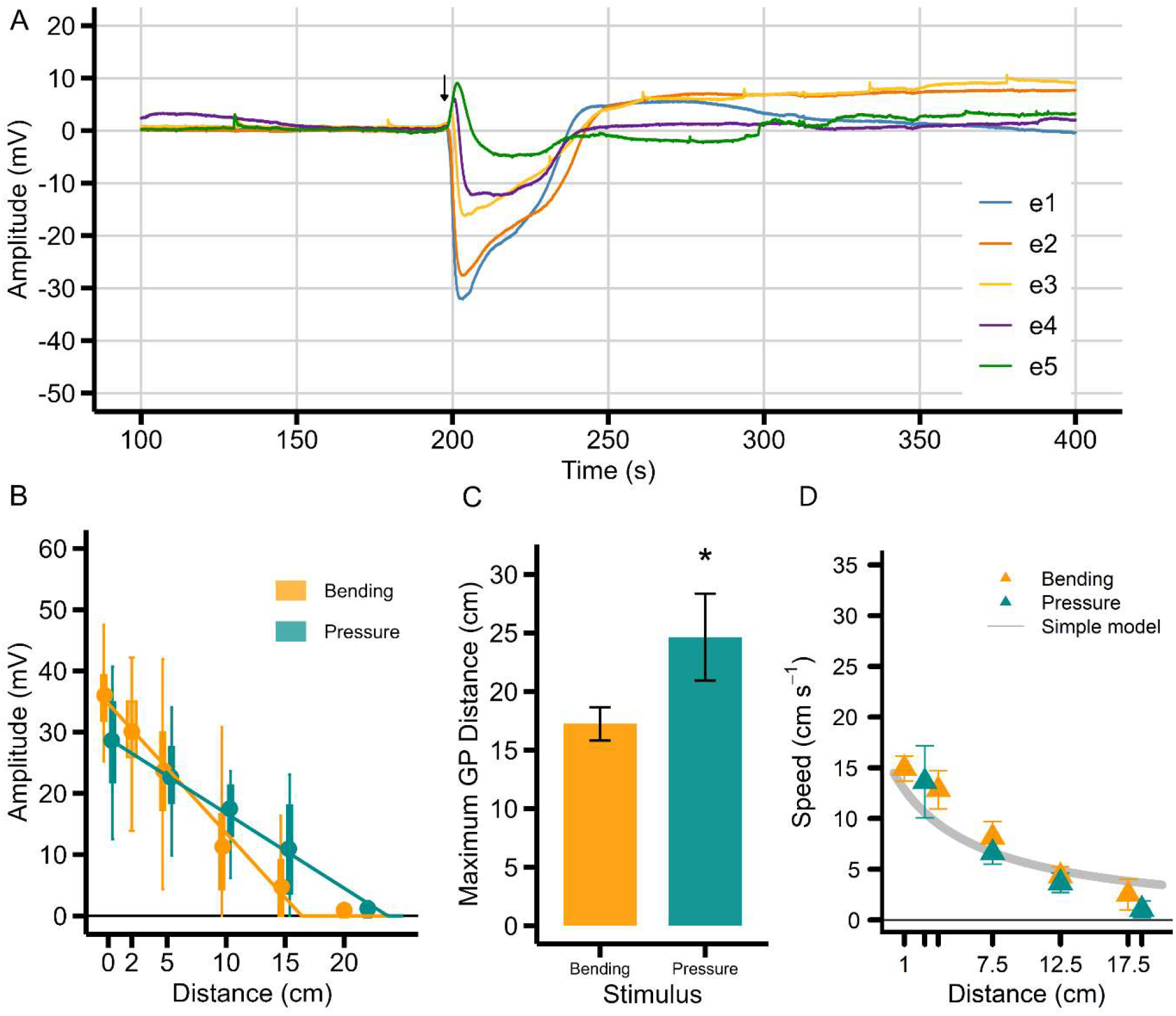
Induction of a depolarization wave through the poplar stem by applying pressure to the root system. (A) Representative 5-channel recording illustrating the depolarization wave induced by applying pressure (0,1 MPa over 10 seconds, indicated by black arrow) to the root system. Electrodes e1 to e5 represent measurements at different distances from the pressure site, as described in Figure 1. (B) Boxplot of the GP’s amplitudes as a function of distance, induced by stem bending (N=40) and by applying pressure to the poplar root system (N=10). Circles indicate the mean. (C) Estimated maximum GP propagation distance (D_max_) in the poplar stem following application of pressure to the root system, compared to stem bending. The asterisk (*) denotes a significant difference compared to bending (95% CI). (D) Mean GP propagation velocity as a function of distance (triangles indicate the mean, error bars represent standard error) for bending (n=40) and pressure to the root system (n=10), overlaid with prediction of simple hydraulic diffusive model 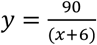 (Simple model, grey line, see Discussion and Supplementary Text S1).

The mean depolarization amplitude (GP) attenuated almost linearly with distance, becoming negligible at 22 cm and never reaching 30 cm (Fig. 6B). Compared to mechanical bending (N=40), the amplitudes induced by root pressurization (N=10) were generally lower but followed a similar attenuation pattern (e.g., at 0 cm: 36.4 ± 2.3 mV for bending vs 28.4 ± 2.9 mV for pressure; at 10 cm: 12.1 ± 2.1 mV for bending vs 9.6 ± 2.7 mV for pressure).

The maximum propagation distance (D_max_) of the depolarization wave confirmed these observations (Fig. 6C). The D_max_ was 17.2 ± 1.4 cm after bending, whereas D_max_ was estimated at 24.7 ± 3.7 cm after root pressurization. This distance for pressure significantly exceeded that observed during bending (*indicated by non-overlapping 95% CI).

Mean GP propagation velocity (Fig. 6D) consistently decreased hyperbolically with distance. Pairwise Wilcoxon rank-sum tests (Holm correction) comparing propagation velocities between the pressure and bending conditions at different distance intervals revealed no statistically significant differences in the mean propagation velocities across the measured distances (all p_adj_ > 0.05). Propagation velocity for pressure ranged from 15.3 ± 4.1 cm·s^−1^ (0-5 cm interval) to 1.2 ± 0.9 cm·s^−1^ (15-22 cm interval), while the fastest propagation for bending was 18.3 ± 2.6 cm·s^−1^ at 1 cm.

These results indicate that hydraulic pressure can initiate GPs with comparable characteristics to those induced by mechanical bending, particularly regarding propagation velocity and pattern of attenuation, but with a significantly longer propagation distance. This supports the hypothesis that propagation length depends on the stimulus.

## Discussion

Long-distance information transfer in trees, crucial for whole-plant acclimation to localized environmental stimuli such as wind-induced bending, remains poorly understood. Previous studies have identified a gradual potential (GP) in poplar and Douglas-fir (Tinturier *et al*., 2021, 2024) triggered by stem bending, distinct from slow wave potentials associated with wounding (Farmer et al., 2020) and action potentials elicited by other stimuli (Lautner et al., 2005; Król et al., 2010). While electrical signals are recognized as early events in plant stress perception (Li *et al*., 2021) and mediators of rapid information propagation, the physical basis for GP generation remains an open question. To address this, we used a biomechanical approach to characterize the relationship between applied deformation intensity and GP characteristics.

### Fundamental Characteristics and Dynamics of GPs

Our results enlighten that the initial amplitude of the GP recorded near the stimulation site remained strikingly consistent (∼35 mV) whatever the intensity of the mechanical stimulation (ε) and the strain rate 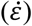. This apparent amplitude saturation suggests that GP generation may operate via an all-or-nothing mechanism, reminiscent of plant action potentials (APs).

Despite this apparent similarity with APs in their generation, GPs diverge notably in their propagation dynamics. While APs maintain constant amplitude over distance (Król *et al*., 2010), GPs exhibit a gradual attenuation, with the rate of decay depending on the stimulus intensity and the strain rate. Specifically, under weaker or slower bending, GPs attenuate more rapidly and propagate over shorter distances. This behavior is distinct from APs, but also differs from electrotonic potentials (EPs), which both attenuate quickly and exhibit initial amplitudes that scale proportionally with stimulus intensity (Volkov et al., 2013; Volkov & Shtessel, 2016) —a feature not observed in GPs.

Our results indicate that the GP amplitude at the stimulation site does not vary significantly with mechanical input strength, distinguishing it from EPs. However, the graded propagation range, modulated by mechanical parameters, and the hyperbolic decrease in propagation velocity suggest a unique form of signal transmission, possibly shaped by hydraulic dissipation or local tissue impedance.

Taken together, GPs appear to occupy a hybrid space between canonical APs and EPs: they may be generated in a binary manner like APs, yet propagate in a graded, intensity-dependent fashion more akin to EPs or SWPs. This duality suggests that GPs may represent a novel class of long-distance electrical signals in plants, particularly suited to encoding both the occurrence and intensity of mechanical stimuli over space and time.

While the initiation and local amplitude of GPs resemble canonical action potentials, their graded propagation and intensity-dependent spatial extent remain unexplained by existing electrical models. These observations raise the question: could GPs be shaped by an interplay between electrical excitability and hydraulic dynamics within the tissue? The following section explores this hypothesis by examining how hydraulic signals might influence GP response or co-propagate with them.

### Hydroelectric coupling model

The observation that both mechanical bending and direct root pressurization generate similar GP responses suggests a mechanistic coupling between hydraulic and electrical signaling. In plants, pressure waves can travel through various tissue compartments—including xylem vessels, apoplast, and symplast—*via* distinct regimes such as inertial or viscous flow (Molz & Ikenberry, 1974; Rockwell et al., 2014). We hypothesize that mechanical stress induces a transient pressure signal, which may subsequently trigger an electrical response through mechanosensitive or hydroresponsive pathways.

Assuming strong coupling between xylem vessel pressure and adjacent parenchyma cells, such a pressure signal could propagate diffusively, with an effective diffusion coefficient D_eff_ [unit m^2^ s^-1^] reflecting the interplay of vascular conductance and tissue elasticity according to the following equation (Supplementary Text S1):

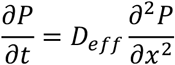

Such a model predicts that pressure amplitude decreases with distance (P_max_ *∼* 1/*x*), and that the apparent propagation velocity decreases hyperbolically *(v ∼* 1/*x*), a behavior strikingly similar to our experimental measurements of GP attenuation and velocity gradients (Figures 2D, 3C, 4D, 5D, 6D).

However, despite these similarities, GPs differ from purely hydraulic waves. A key difference lies in their amplitude behavior: diffusion theory predicts that hydraulic signal amplitude should be proportional to the stimulus intensity, while we observe that GP amplitude remains almost constant across different bending deformations (Figure 2B) and bending strain rate (Figure 3A). Additionally, the temporal profile of GP signals does not align with that of a diffusion process, as GP signals exhibit a narrowing width with distance (Figures 2A, 4A, 5A), while a diffusive wavefront would be expected to spread.

A notable result is that GP depolarization is followed by a recovery phase in both bending (transient pressure) and pressurization experiments (constant applied pressure) (Figures 2A, 6A). This suggests that the relevant trigger for GP generation may not be the absolute value of the pressure, but rather its time derivative or another non-linear transduction mechanism. If so, GPs would not result directly from hydraulic wave propagation, but rather represent an active cellular response to pressure variations. Further experiments comparing gradual and abrupt pressure changes could help to determine whether the GP response is governed by the rate of pressure rather than by its absolute value.

Taken together, these observations are difficult to reconcile with a simple, direct conversion of a hydraulic pressure wave into an electrical GP signal. Instead, our results suggest that GP signals emerge from a non-linear transduction pathway that couples mechanical stress, hydraulic fluctuations, and electrical activity. Alternatively, GP propagation may be electrically autonomous, modulated by hydraulic conditions but not directly driven by them. The observed symmetry in pressure variations during bending and unbending (Lopez et al., 2014; Louf et al., 2017), combined with the similarity in GP characteristics, reinforces the hypothesis that plants are capable of detecting and responding to mechanical state changes in both directions. This non-directional sensing could provide plants with a more nuanced perception of dynamic mechanical environments.

Future research should focus on simultaneous measurements of pressure and electrical signals in response to the same stimulus, to determine whether pressure variations directly modulate GP generation, or if GP propagation follows an independent electrical pathway influenced by hydraulic state changes.

### Physiological Significance and Adaptive Value of GPs

Beyond the biophysical properties of GPs and their potential hydraulic coupling, their consistent occurrence under various mechanical contexts raises an important question: what is their physiological function and adaptive significance? Here, we examine how GPs reflect and possibly contribute to the plant’s dynamic perception of mechanical states.

### Bidirectional sensitivity and mechanical state changes

Our experiments reveal that GPs with similar characteristics are generated both during initial stem bending and upon release after 4 hours of sustained deformation (Figure 5). This observation aligns with the transient nature of local mechanical responses observed across different scales and systems. At the cellular level, mechanosensitive channels exhibit rapid activation followed by quick inactivation under sustained membrane tension in patch-clamp experiments (Peyronnet et al., 2014). Similarly, pressure measurements in bent stems show a rapid peak that decreases while the stem is still under deformation (Lopez *et al*., 2014). This transient response pattern is also evident in calcium signaling: as demonstrated by Shih *et al*. (2014) who showed that cellular calcium peaks diminish before the mechanical stress is released in *Arabidopsis* roots.

This bidirectional sensitivity mirrors calcium responses in *Arabidopsis* roots, where Shih et al. (2014) reported calcium signals triggered by both stretching and release. Notably, one of these calcium peaks exhibited refractory periods, reminiscent of the GP characteristics we observed. Furthermore, the involvement of the FERONIA receptor in these calcium responses, coupled with the overexpression of FERONIA homologs in bent poplar stems (Pomiès et al., 2017), suggests potential conservation of mechanosensing mechanisms across plant species and organs.

### Temporal integration through refractory period

Our results reveal a temporally structured refractory response in GP dynamics following mechanical stimulation. When a second bending stimulus was applied 2.5 minutes after the initial flexion, we observed a significant reduction in both GP amplitude and propagation distance, suggesting a transient decrease in tissue excitability (Figure 4).

Interestingly, the response to a second bending applied after a 4 hours delay shows a different profile: the GP amplitude near the bending zone was fully restored, indicating that local excitability had returned to baseline. But the signal propagated over a significantly shorter distance compared to the initial response. This suggests that, while the capacity for local signal generation was reestablished, long-distance transmission remained impaired.

After a delay of 7 hours after the first stimulation, both GP amplitude and propagation distance were comparable to those observed during the initial response, indicating a full recovery of the signaling system.

While refractory periods are well characterized in plant action potentials (APs), where they limit the frequency of signal generation (Paszewski & Zawadzki, 1976; Favre & Agosti, 2007), the dynamics observed here in GPs differ notably. In addition to the short-term reduction in excitability seen after 2.5 minutes, we identify a second, longer-lasting effect after 4 hours that selectively affects signal propagation. This pattern suggests a multi-phase regulation of signal transmission, beyond the simple inactivation of ion channels.

The restoration of full amplitude after 4 hours implies that the generation of the signal itself is not impaired. Instead, the reduced propagation range may reflect transient alterations in the tissue’s conductive properties. Possible contributing factors include local ionic imbalances, hydraulic adjustments, or temporary changes in tissue permeability that affect signal diffusion.

Another hypothesis relates to mechanical adaptation at the cellular level. As proposed by Audemar *et al*. (2023), repeated mechanical stimulation can increase cellular rigidity, thereby limiting membrane stretch and the activation of mechanosensitive channels. Such mechanical stiffening may be more pronounced in distal tissues, restricting GP propagation while leaving local generation unaffected. The progressive recovery observed after 7 hours could reflect a reversible relaxation of this mechanical state, allowing normal signal transmission to resume.

This temporary reduction in signal transmission efficiency appears to be part of a broader mechanical acclimation process. Martin *et al*. (2010) reported a longer-term desensitization in poplar stems, where gene regulatory responses decreased after repeated bending and persisted for up to five days. Together, these findings support a hierarchical model of mechanical memory in plants, where rapid electrical adjustments (from minutes to hours) precede and possibly facilitate longer-term molecular reprogramming (days scale), enabling dynamic tuning of the plant’s sensitivity to mechanical stress.

### Adaptive significance

The observed modulation of GP propagation distance by the velocity and intensity of stem flexion provides a compelling mechanism for plants to spatially discriminate mechanical stimuli mimicking wind forces. While the consistent maximal GP amplitude near the point of stimulation suggests a reliable initial detection of mechanical perturbation, the subsequent attenuation and limited propagation of GPs induced by weaker or slower flexions (mimick gentle breezes or minor deflections) may reflect a “signal filtering.” This spatially restricted signaling could prevent the allocation of resources for systemic responses to transient or low-impact mechanical stimuli that do not pose a significant structural threat. Conversely, the more extensive propagation of GPs elicited by stronger or faster flexions (analogous to stronger gusts or rapid movements) could trigger more widespread physiological and developmental adjustments necessary for acclimation to significant mechanical stress.

This stimulus-dependent spatial encoding, coupled with the temporal filtering provided by the refractory period, suggests a sophisticated system for integrating complex mechanical information. The bidirectional sensitivity, allowing the detection of both stem displacement and its release, further enhances the plant’s ability to track dynamic wind patterns. By not only sensing the occurrence but also gauging the intensity and rate of change of wind-induced mechanical stress through the spatial reach of the GP, and by temporally filtering repetitive stimuli, the plant can optimize resource allocation for adaptive growth responses. For instance, prioritizing responses to sustained high winds over brief, low-intensity gusts would be energetically advantageous. This finely tuned electrophysiological response to mechanical cues underscores that plant mechanosensing is an active process where stimulus characteristics are transduced into spatially and temporally modulated signals, enabling a nuanced perception of the dynamic mechanical environment crucial for survival and adaptation, particularly in wind-exposed habitats.

## Conclusion and perspectives

Our findings highlight a complex mechanosensory system in poplar, in which GP integrate mechanical cues across both spatial and temporal scales. This electrical response to bending displays characteristic refractory dynamics that modulate signal propagation, and points toward a possible coupling between hydraulic and electrical phenomena. Root pressurization alone triggers a GP in the stem, even in the absence of bending. This observation supports the hypothesis of hydroelectric coupling, where pressure fluctuations in the vascular system may directly elicit electrical signals. The similarity in propagation speeds between GPs and hydraulic pulses reinforces this hypothesis, although the strong, nonlinear attenuation of GP amplitude suggests the presence of additional regulatory layers. These results support the view that vascular system — as hinted by comparative data from Douglas fir and poplar (Tinturier et al., 2024) — plays a central role in shaping signal transport.

A key question is whether GPs act as functional biological signals — capable of influencing downstream processes such as gene expression, hormonal pathways, or growth regulation. Identifying the molecular mechanisms responsible for GP generation and propagation will be essential to place this process within broader plant mechanotransduction networks.

These insights have potential agronomic implications. Engineering plants with enhanced mechanosensory properties could improve resistance to wind stress, lodging, or mechanical damage. Moreover, understanding how vascular mechanics regulate signal propagation could open new avenues for tuning long-distance signaling in crops.

In conclusion, this study provides new evidence for an hydroelectric mechanism in plant mechanosensing. Demonstrating that pressure fluctuations alone can elicit electrical responses, it raises the possibility that GP may serve as integral components of whole-plant signaling networks. Unraveling their functional role in systemic acclimation remains a central challenge for future research.

## Abbreviations

AP: Action Potential
CLE25: CLAVATA3/EMBRYO SURROUNDING REGION-related 25
D_eff_: Effective diffusion coefficient
D_max_: Maximum propagation distance
EP: Electrotonic Potential
GP: Gradual Potential
L_0_: Length of bent segment (12 cm)
Lb: Length of bent segment
miRNA: microRNA
ROS: Reactive Oxygen Species
S3M: Strain Sensing and Signaling Model
SE: Standard Error
SWP: Slow Wave Potential
ε_max_: Maximum peripheral longitudinal strain
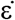: Strain rate
ρ: Radius of curvature

## Supplementatry data

The following supplementary data are available at JXB online.

Text S1. Physical model of hydraulic signal propagation with diffusion equations and parameter estimations.

## Acknowledgments

The authors thank Amélie Coston for the poplar production; Stéphane Ploquin and Têtè Sévérien Barigah for their technical support.

## Author contributions

JL.J, E.B, and E.T conceived the original screening and research plans. JL.J, E.B, N.L.B, and E.T designed the experiments. The experimental work was carried out and interpreted by E.T, JL.J, and E.B. GG and YF developed and wrote the hydroelectric model. E.T wrote the first draft of the article. All authors contributed to the writing and revision of the article.

## Data availability statement

The data that support the findings of this study are available from the corresponding author upon reasonable request.

